# Hippocampus Glutathione S Reductase Potentially Confers Genetic Resilience to Cognitive Decline in the AD-BXD Mouse Population

**DOI:** 10.1101/2024.01.09.574219

**Authors:** Michael C. Saul, Elizabeth M. Litkowski, Niran Hadad, Amy R. Dunn, Stephanie M. Boas, Jon A. L. Wilcox, Julia E. Robbins, Yiyang Wu, Vivek M. Philip, Gennifer E. Merrihew, Jea Park, Philip L. De Jager, Dave E. Bridges, Vilas Menon, David A. Bennett, Timothy J. Hohman, Michael J. MacCoss, Catherine C. Kaczorowski

## Abstract

Alzheimer’s disease (AD) is a prevalent and costly age-related dementia. Heritable factors account for 58-79% of variation in late-onset AD, but substantial variation remains in age-of-onset, disease severity, and whether those with high-risk genotypes acquire AD. To emulate the diversity of human populations, we utilized the AD-BXD mouse panel. This genetically diverse resource combines AD genotypes with multiple BXD strains to discover new genetic drivers of AD resilience. Comparing AD-BXD carriers to noncarrier littermates, we computed a novel quantitative metric for resilience to cognitive decline in the AD-BXDs. Our quantitative AD resilience trait was heritable and genetic mapping identified a locus on chr8 associated with resilience to AD mutations that resulted in amyloid brain pathology. Using a hippocampus proteomics dataset, we nominated the mitochondrial glutathione S reductase protein (GR or GSHR) as a resilience factor, finding that the DBA/2J genotype was associated with substantially higher GR abundance. By mapping protein QTLs (pQTLs), we identified synaptic organization and mitochondrial proteins coregulated in trans with a cis-pQTL for GR. We found four coexpression modules correlated with the quantitative resilience score in aged 5XFAD mice using paracliques, which were related to cell structure, protein folding, and postsynaptic densities. Finally, we found significant positive associations between human *GSR* transcript abundance in the brain and better outcomes on AD-related cognitive and pathology traits in the Religious Orders Study/Memory and Aging project (ROSMAP). Taken together, these data support a framework for resilience in which neuronal antioxidant pathway activity provides for stability of synapses within the hippocampus.

## Introduction

Alzheimer’s disease (AD) is a prevalent and costly age-related dementia. By 2050, an estimated 14 million Americans will have AD with an associated healthcare cost of approx. $1.1 trillion^1^.

Genetics is a key determinant of AD risk; heritable factors account for 58-79% of variation in late-onset AD^2^. However, despite its strong genetic etiology, there is substantial variation in age-of-onset^3^, disease severity^4^, and whether those with high-risk genotypes acquire AD^5^. Genetic factors appear to confer resilience against even strongly penetrant autosomal dominant AD risk loci^5–7^. Our understanding of the biological systems governing cognitive resilience to AD is emerging^8^. The genetic origins of resilience, though less tractably studied with genetics than AD risk itself, are of great interest as they may reveal therapeutic targets to ameliorate clinical outcomes among those with genetic predisposition for AD^8^.

The AD-BXD mouse population is a powerful emerging model for studying AD resilience. AD-BXD mice query two different levels of genetic information: strains and genotypes^9^. The genetically diverse BXD strains are a mosaic of the approximately 6 million segregating single nucleotide polymorphisms between C57BL/6J and DBA/2J mice^9–12^. The AD genotype consists of a transgene bearing five pathogenic variants in two human genes (5XFAD) ^13^: human *APP* carrying three pathogenic variants and human *PSEN1* carrying two pathogenic variants. 5XFAD animals can be contrasted with non-transgenic strain-matched control animals (Ntg).

Reproducible BXD strains allow repeated measurement of the same mosaic genomes^9^, a strategy that boosts the mappable heritability of genetic loci^14^. This property facilitates quantitative trait locus (QTL) mapping of the relatively lower effect size variants that modify susceptibility to Alzheimer’s-like cognitive phenotypes in 5XFAD animals. Further, the same cognitive phenotype can be measured in both 5XFAD and Ntg mice within the same background strain. The ability to contrast a strain’s performance with different AD genotypes allows the measurement of novel and interesting derived phenotypes based upon the property of trait correlation^9,12^. Because of these properties, our lab has successfully leveraged AD-BXD mice to identify multiple genetic modifiers of AD-related phenotypes in recent years^6,9,15,16^.

Much of our past genomics work on AD-BXD mice has utilized transcript abundance as a proxy for gene expression. This strategy has successfully identified transcriptional regulatory network features that predict AD resilience^17,18^. However, protein abundance is a substantially stronger predictor of AD pathophysiology than RNA abundance^19,20^. Data-independent acquisition (DIA) proteomics is a powerful proteomics technology for the measurement of proteins and peptides from complex mixtures^21^. With the use of a DIA-based chromatogram library^22,23^, DIA proteomics improves the measurement of peptides across a large number of samples. The addition of quantitative proteomics to AD-BXD genetics studies represents a significant advance over previous methodologies for gene expression genetics^24^, enabling one to map protein quantitative trait loci (pQTL) for thousands of proteins.

Here, we quantified resilience to cognitive decline using a novel regression-based metric that compares each strain’s cognitive performance with an AD transgene to what its cognitive performance would be like without the AD transgene. We utilized this resilience quantitative trait measure to map individual strain differences in resilience to the AD transgene to a genetic locus. Given that the locus harbored a number of potential candidate genes, we then leveraged proteomics data to nominate mitochondrial glutathione S reductase (GR or GSHR) as the likely molecular genetic mechanism for resilience. GR is a key component of the endogenous antioxidant system, and functions to recycle oxidized glutathione using nicotinamide adenine dinucleotide phosphate (NADPH). We found potential novel targets for AD by querying elements coregulated in trans with GR. Finally, we implicated differential expression of networked synaptic proteins as the likeliest downstream consequences of differences in resilience.

## Results

### Deriving and Mapping a Quantitative Trait for Cognitive Resilience to FAD Mutations

Our previous work has demonstrated the ability of the AD-BXD mouse panel to yield insights about the genetic basis of variation in AD phenotypes^9^ and to identify potential new biological drivers of resilience^25^. Though this work demonstrated the clear value of the AD-BXD mouse panel in identifying resilience factors, substantial unexplained variance in resilience has remained inaccessible to our previous methodologies. Here we utilized genetics tools available to us with the AD-BXD mouse panel to map resilience as a continuously varying quantitative trait.

We operationalized resilience as an individual’s cognitive function being relatively intact compared to expectations. We defined cognitive function using the behavioral measure of percent time freezing in a contextual fear memory task (CFM). We measured CFM in both 5XFAD and Ntg at both 6 months and 14 months of age. To assess how well resilience could be measured using this approach, we regressed 14-month 5XFAD values for CFM against the strain mean of 14-month Ntg CFM values from the same strain (male and female samples pooled), weighting the regression by 1/*n*_*BXD*_ where *n*_*BXD*_ is the number of 5XFAD samples per BXD strain. As CFM is a ratio, CFM values were arcsine transformed for regression, though results were back transformed for plotting and interpretation. The result is a regression line with identical slope and intercept to a regression of BXD strain means, but with an ability to derive individual-level residuals for the 5XFAD mice.

In the regression of 14mo CFM for 5XFAD mice on strain means of 14mo CFM for Ntg mice of the same BXD background, we found that a statistically significant but small amount of variance was explained (R^2^ = 0.091, p = 0.001) and that the regression line had a slope less than 1 (approx. 0.48, **Figure 1A**). This result indicates that the 5XFAD transgene generally decreases cognitive function, but a substantial amount of variance in the cognitive decline experienced by an individual 5XFAD animal is not explained by its relationship with an Ntg animal from the same strain. To measure this additional unexplained variance, we calculated the standardized residuals of the linear regression. We refer to this value as the quantitative resilience trait, which varies across the AD-BXD strains and is heritable (**Figure 1B**, ℎ^$^ = 0.516, ℎ^$^ _̅_ = 0.83), indicating that approximately 50% of the variance in cognitive resilience to AD carrier status is governed by genetic factors.

**Figure 1:**
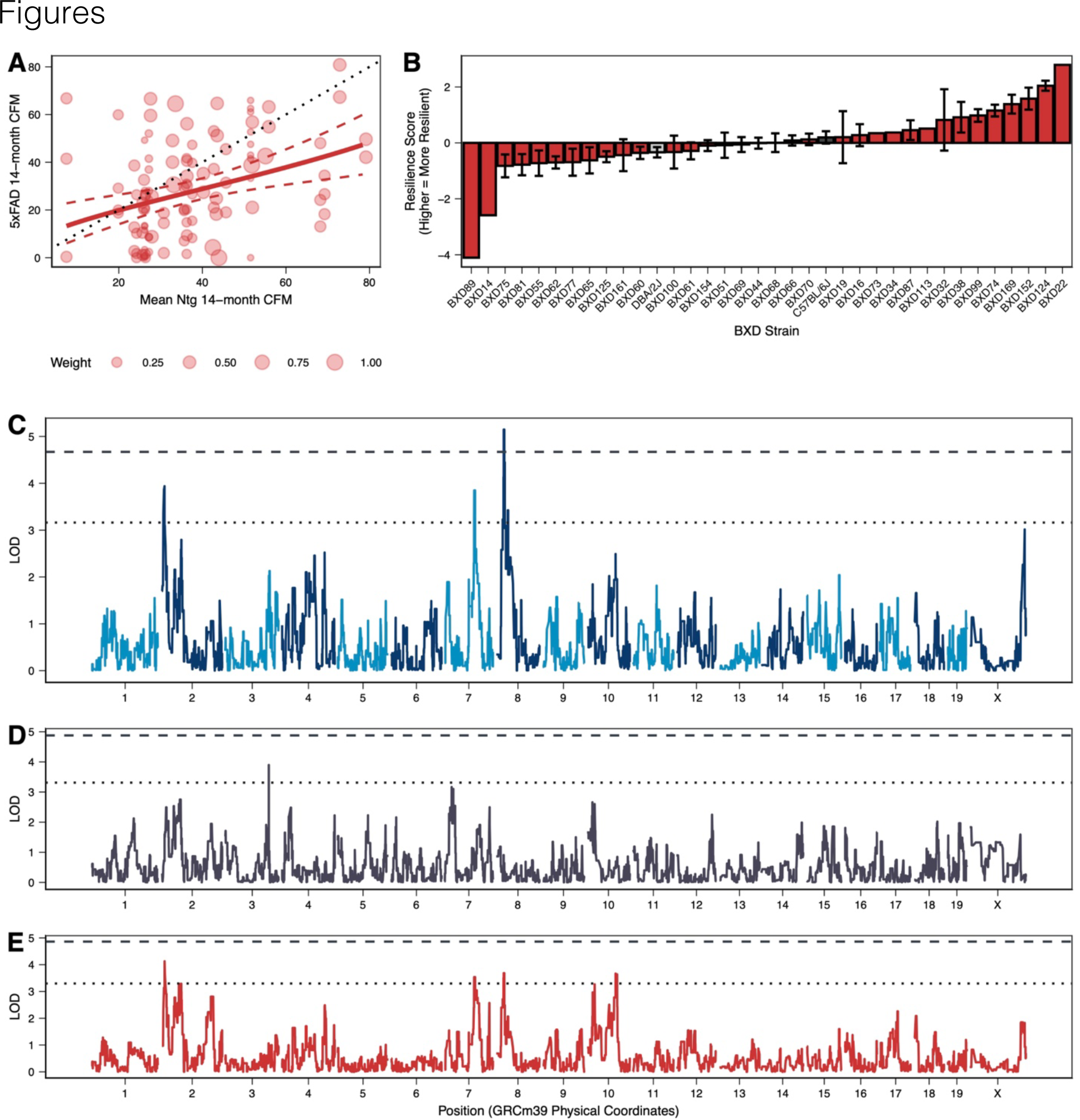
Resilience to AD-related cognitive decline is quantitative, heritable, and maps to chr8. **A)** Regression analysis of all 14-month 5XFAD contextual fear memory observations against strain means of 14-month Ntg contextual fear memory (R^2^ = 0.09, p = 0.0014). Observations are weighted by n^-1^ to ensure equal weight is given to each BXD strain. **B)** Strain survey of resilience scores, calculated from normalized residuals of the regression analysis (ℎ^$^ = 0.516, ℎ^$^ _̅_ = 0.83). **C)** Mapping results for rank-normal transformed resilience scores reveal an FDR < 0.05 QTL on chr8. Dashed line is a permutation-based threshold of 0.05; dotted line is the suggestive QTL threshold of 0.63. **D)** Mapping results of contextual fear memory in Ntg animals does not show the same QTL on chr8. **E)** Mapping results of contextual fear memory in 5XFAD animals does not show the same QTL on chr8.

We next sought to understand the specific genetic mechanisms governing the strain differences across the AD-BXD panel. Using a total of 34 AD-BXD strains bearing the 5XFAD transgenes, we mapped the strain means of resilience. This approach identified a strong quantitative trait locus (QTL) at the FDR < 0.01 level on chr8 at approximately 25.87 Mb (**Figure 1C**, peak empirical FDR = 0.008, 99% Bayes credible interval: chr8:18.55-38.46 Mb in GRCm39/mm39 coordinates). This result is unique to the quantitative resilience trait; QTL mapping Ntg CFM does not return a peak on chr8 (**Figure 1D)**, and mapping 5XFAD CFM returns a relatively small chr8 peak at the suggestive level (**Figure 1E**). Together, these results indicate that the chr8 locus modifies the degree to which animals display cognitive resilience to AD mutations.

### Integrating Proteomics Implicates Mitochondrial Glutathione S Reductase in Resilience

Because the AD-BXDs are derived from an F1 breeding of a 5XFAD mouse on a C57BL/6J backcross background and a BXD strain, the two genotypes present in the AD-BXD across the genome are: homozygous for C57BL/6J (B6/B6) and animals heterozygous for a DBA/2J genotype (B6/D2). The allele effect at the peak marker (rs33539160) indicates that the B6/D2 genotype confers resilience relative to B6/B6, implicating the D2 genotype as the resilient genotype (**Figure 2A**). The mapped QTL interval includes many protein-coding genes. We used SnpSift to identify 35 DBA/2J nsSNPs predicted to be deleterious in 23 protein-coding genes within the Bayes credible interval (**Supplementary Table S1**). Of these SNPs, only 4 were in 3 genes whose proteins had measurable abundances within our DIA proteomics dataset: *Prag1* (rs47798524 and rs30475069), *Pomk* (rs32817877), and *Agpat5* (rs3546362755). After a review of the consequences of these 4 nsSNPs (brief summaries in **Supplementary Table S1**), there were no obvious cases in which these nsSNPs explained increased resilience in the DBA/2J genotype relative to the C57BL/6J genotype, though we cannot completely preclude the possibility of an nsSNP as a genetic driver of the resilience phenotype.

**Figure 2:**
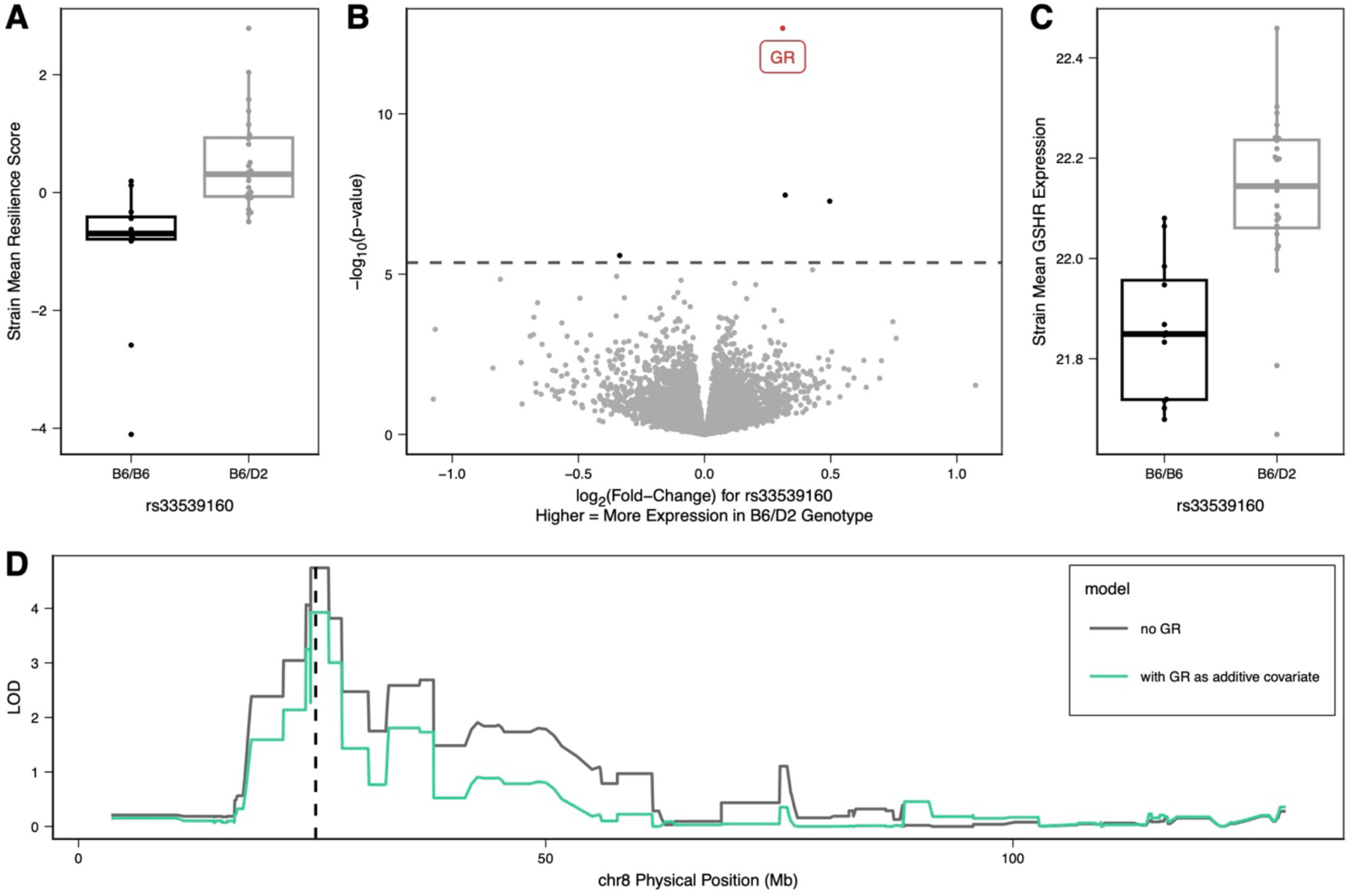
Analysis of proteomics data reveals mitochondrial glutathione-disulfide reductase (GR) as potential factor within the QTL interval. **A)** Strain mean resilience scores of different 5XFAD AD-BXD strains stratified by genotype at rs33539160, the peak chr8 marker. At the peak marker, the B6/D2 heterozygote genotype shows higher resilience than the B6/B6 homozygote genotype. **B)** Differential protein expression by stratifying by genotype at rs33539160 reveals the protein GR as the strongest finding. GR is encoded by the gene *Gsr*, which lies entirely within the 99% Bayes credible interval for the chr8 resilience QTL. Dashed horizontal line corresponds to q-value of 0.01. **C)** Strain mean GR expression of different 5XFAD AD-BXD strains stratified by genotype at rs33539160, the peak chr8 marker. The allele expression pattern of GR mirrors that seen in the resilience plot. Higher expression of GR is associated with higher resilience. **D)** A mapping model with GR as an additive covariate reduces the LOD score at the peak marker (LOD difference = 0.82, at rs33539160, vertical dashed line), which provides some evidence that GR protein abundance mediates the genotype effect at this marker.

To nominate potential protein abundance effectors of this phenotype, we stratified DIA proteomics data for 14-month 5XFAD animals by their genotype at rs33539160, the top marker on chr8. We used a conservative q-value threshold of 0.01 to ensure that findings were highly likely to be related to the top marker. Of the 8,591 proteins detected in the protein samples, 4 were differentially abundant at q < 0.01 according to their rs33539160 genotype (**Figure 2B**, **Supplementary Table S2**). The top protein was mitochondrial glutathione-disulfide reductase (GR, UniProt P47791, q = 1.53x10^-9^). The gene *Gsr* encodes this protein and lies on chr8 positioned at approximately 34.14-34.19 Mb, completely within the 99% Bayes credible interval for the chr8 QTL. GR was the only protein of the 4 differentially abundant proteins encoded by a gene on chr8. The allele effect at rs33539160 for GR was similar to the effect for resilience, indicating that increased protein abundance in DBA/2J heterozygote mice relative to C57/BL6 homozygotes is the potential explanatory factor for resilience (**Figure 2C**). Mediation analysis with GR protein abundance at rs33539160 trends toward providing evidence of GR as a mediator, though it fails to reach statistical significance (ACME p-value = 0.084, **Figure 2D**).

The mediation analysis constitutes modest evidence of a potential causal effect, one that may be diluted by a difference in timing between the causes and the outcome of increased resilience.

Together, these results nominate variation in GR protein abundance as a potential factor in increased cognitive resilience among AD-BXD strains bearing the DBA/2J allele.

### Protein QTL Analysis Identifies Related Molecular Processes in the GR Mechanism

The GR protein catalyzes the reduction of oxidized glutathione (GSSG) into reduced glutathione (GSH) driven by NADPH. An increased abundance of GR may affect cellular response to oxidative stress through multiple mechanisms. Increased ability to resist oxidative stress may be favorable to the survival of cells in the nervous system that are subject to high degrees of oxidative damage. Oxidative stress is believed to be highly important to the pathophysiology of AD^26^.

To expand the GR mechanism, we utilized protein QTL (pQTL) data, which allow us to query how genetic mechanisms act in trans to affect systems of molecules. Performing a pQTL analysis on 14-month 5XFAD mice, there was a significant cis-pQTL for GR on chr8 (**Figure 3**).

**Figure 3:**
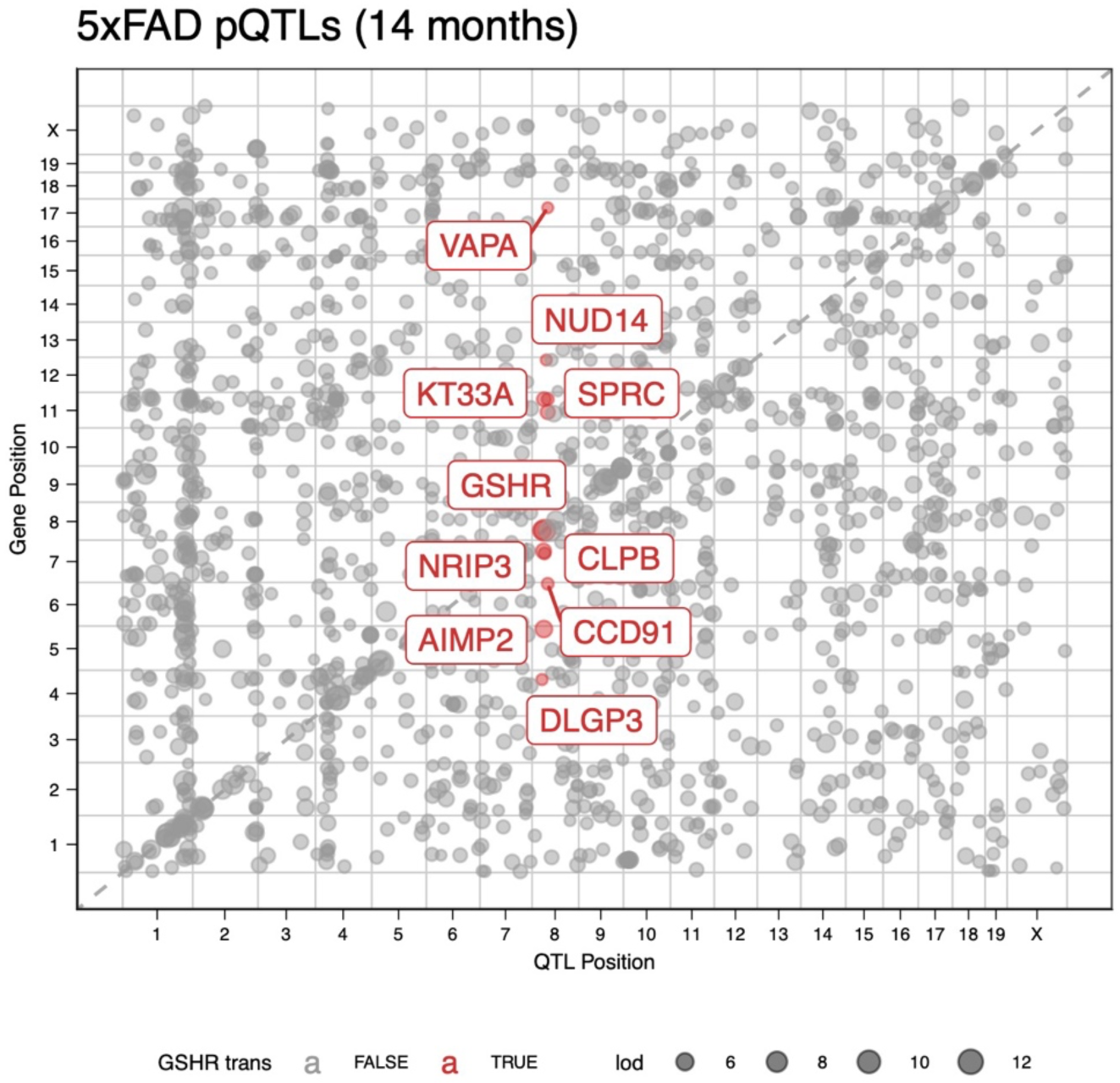
Trans-pQTL plot highlights proteins coregulated in trans with the GR cis-pQTL on chr8. These proteins include DLGP3, which has been studied in the context of schizophrenia and obsessive-compulsive disorder; the mitochondrial disaggregase CLPB; the aminoacyl tRNA synthase complex interacting protein AIMP2; the extracellular matrix regulatory protein SPRC; the keratin protein KT33A; the nuclear receptor interacting protein NRIP3; the coiled-coil domain containing protein CCD91; and the vesicle-associated membrane protein-associated protein VAPA.

In addition, we found that approximately 9 proteins were significantly coregulated in trans (on a different chromosome or farther than 15 Mb from the GR peak) within 15 Mb of the cis-pQTL for GR at a LOD score > 4 (**Figure 3**). These proteins were diverse and included the protein DLGP3, a neuronal scaffolding protein involved in postsynaptic density formation. DLGAP proteins have been associated with a wide variety of neurodegenerative diseases^27^, and the closely related gene *Dlgap2* was identified as a genetic regulator of age-related cognitive decline in Diversity Outbred mice in our lab^28^. Based upon these results, we speculate that GR protein level differences between B6/B6 and B6/D2 genotypes confer resilience by making neurons more tolerant of oxidative stress, potentially leading to higher stability of postsynaptic densities.

### Proteome Coexpression Reveals a Tightly Clustered Protein Coexpression Network of Resilience

To identify the biological systems most strongly associated with differences in cognitive resilience to AD mutations, we used the paraclique method^29^ to construct small and tightly coregulated coexpression clusters *de novo* in 14-month aged 5XFAD proteomics samples. This method constructed a total of 21 paracliques containing between 10 and 623 proteins each. Of these 21 paracliques, four (Paracliques 9, 16, 20, and 21) had eigengenes with a significant correlation to the quantitative resilience trait (q < 0.10, **Table 1**).

**Table 1:** Paraclique eigengene correlation with resilience. Paracliques 9, 16, 20, and 21 showed significant relationships with resilience.

| paraclique | # proteins | eigengene-resilience bicor | bicor p-value | bicor q-value | significant |
| --- | --- | --- | --- | --- | --- |
| 1 | 10 | 0.04 | 0.70 | 0.29 |  |
| 2 | 11 | 0.07 | 0.48 | 0.26 |  |
| 3 | 12 | 0.07 | 0.50 | 0.26 |  |
| 4 | 13 | 0.13 | 0.17 | 0.22 |  |
| 5 | 17 | 0.07 | 0.44 | 0.26 |  |
| 6 | 18 | 0.07 | 0.45 | 0.26 |  |
| 7 | 22 | -0.08 | 0.39 | 0.26 |  |
| 8 | 26 | -0.07 | 0.44 | 0.26 |  |
| 9 | 38 | 0.20 | 0.04 | 0.10 | * |
| 10 | 50 | -0.08 | 0.40 | 0.26 |  |
| 11 | 53 | -0.02 | 0.84 | 0.33 |  |
| 12 | 53 | -0.05 | 0.64 | 0.29 |  |
| 13 | 62 | -0.15 | 0.13 | 0.20 |  |
| 14 | 68 | 0.04 | 0.70 | 0.29 |  |
| 15 | 68 | 0.00 | 0.97 | 0.36 |  |
| 16 | 74 | 0.29 | 0.00 | 0.02 | * |
| 17 | 95 | -0.11 | 0.23 | 0.24 |  |
| 18 | 104 | 0.04 | 0.69 | 0.29 |  |
| 19 | 155 | -0.11 | 0.25 | 0.24 |  |
| 20 | 299 | -0.19 | 0.05 | 0.10 | * |
| 21 | 623 | -0.20 | 0.04 | 0.10 | * |

To query the biological mechanisms underpinning these paracliques, we tested Gene Ontology Biological Process (GO BP) enrichment within this cluster. GO BP analysis identified many terms per cluster. To reduce the dimensionality and understand the generalized mechanisms within these terms, we analyzed the semantic similarity of GO terms and used multidimensional scaling on the resultant similarity matrix. Visualized similarities generally implicate multiple systems including: cell shape and structure for Paraclique 9, protein folding and chaperones for Paraclique 16, cell shape and transport for Paraclique 20, and biological processes relevant to neurons including postsynaptic organization and regulation of neurotransmitter secretion and neurotransmitter levels for Paraclique 21 (**Figure 4**). Together, these results sketch a diverse set of mechanisms that correlate with resilience. The coexpression data support the hypothesis that resilience operates at least in part by conserving the structures of postsynaptic densities through changes in cell structure, protein folding, and postsynaptic density formation.

**Figure 4:**
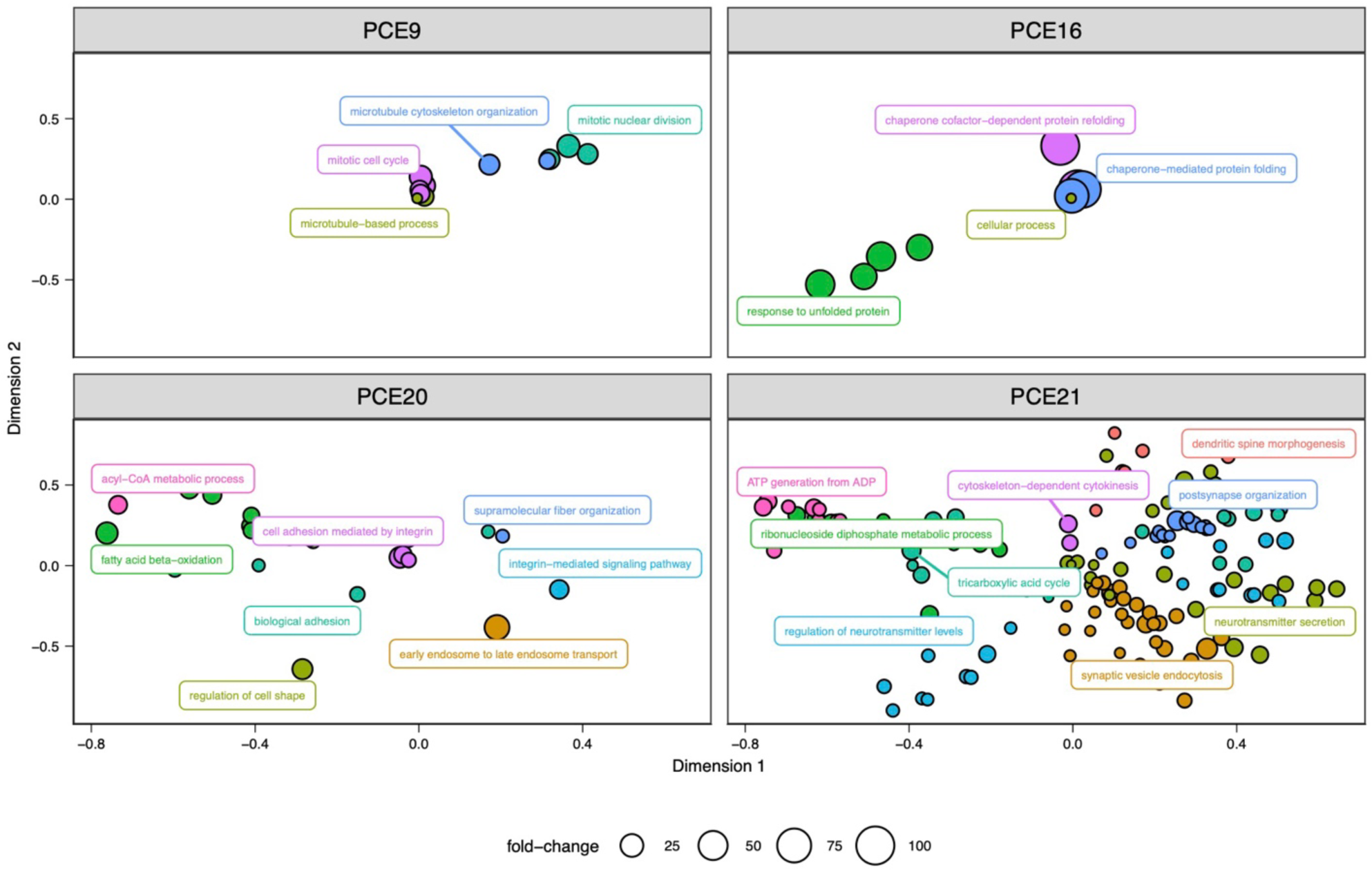
Gene Ontology clustering plot of GO BP terms enriched in Paracliques 9, 16, 20, and 21. These data indicate that the four paracliques that reached significance query multiple and sometimes disparate biological processes. For example, PCE9 mostly includes microtuble and mitosis-related biological processes, PCE16 appears to relate to protein folding, PCE20 includes processes related to cell shape and transport, and PCE21 includes a diversity of biological processes relevant to neurons including postsynaptic organization and regulation of neurotransmitter secretion and neurotransmitter levels.

### Greater *GSR* Transcript Abundance Is Associated with Better Cognition and Less Pathology in Human Samples

The murine gene encoding mitochondrial glutathione S reductase, *Gsr*, is a 1:1 ortholog of the human gene *GSR* (OrthoDB v11, Euarchontoglires group <u>81598at314146</u>). To evaluate the relationship of the human *GSR* gene with cognitive function and AD-related pathology in humans, we first assessed bulk RNA sequencing (RNAseq) data from individuals in the ROSMAP longitudinal cohorts^30,31^. We used linear regression models (for cross-sectional outcomes) and linear mixed-effects models (for longitudinal outcomes) to evaluate the relationship of *GSR* transcript abundance and several outcomes: global cognition at last visit, longitudinal global cognitive trajectories, and AD pathology from autopsy (amyloid and tangle burden). These outcomes were evaluated in three brain regions: the caudate nucleus (CN), the dorsolateral prefrontal cortex (DLPFC), and posterior cingulate cortex (PCC). Demographic information about the populations has been previously published^32^.

Greater *GSR* gene expression was associated with better longitudinal cognitive trajectories and cross-sectional cognitive function in CN, DLPFC, and PCC (**Table 2**). Additionally, greater gene expression was associated with less amyloid beta load, fewer NP, and lower tau tangle density in CN, CLPFC, and PCC (**Table 3**). Together, these results indicate that greater *GSR* expression is associated with the amelioration of AD-related cognitive and pathology phenotypes.

**Table 2:**
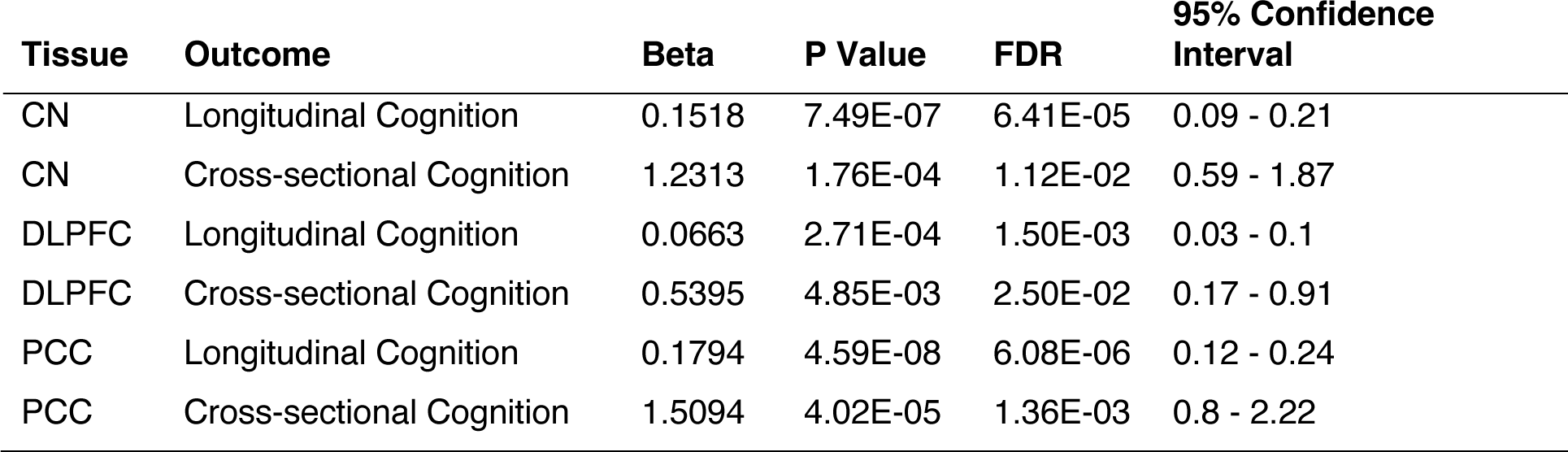
*GSR* Gene Expression Association with Cognition in Humans.

| **Tissue** | **Outcome** | **Beta** | **P Value** | **FDR** | **95% Confidence Interval** |
| --- | --- | --- | --- | --- | --- |
| CN | Longitudinal Cognition | 0.1518 | 7.49E-07 | 6.41E-05 | 0.09 - 0.21 |
| CN | Cross-sectional Cognition | 1.2313 | 1.76E-04 | 1.12E-02 | 0.59 - 1.87 |
| DLPFC | Longitudinal Cognition | 0.0663 | 2.71E-04 | 1.50E-03 | 0.03 - 0.1 |
| DLPFC | Cross-sectional Cognition | 0.5395 | 4.85E-03 | 2.50E-02 | 0.17 - 0.91 |
| PCC | Longitudinal Cognition | 0.1794 | 4.59E-08 | 6.08E-06 | 0.12 - 0.24 |
| PCC | Cross-sectional Cognition | 1.5094 | 4.02E-05 | 1.36E-03 | 0.8 - 2.22 |

**Table 3:** *GSR* Gene Expression Association with Neurodegenerative Pathology in Humans.

| Tissue | Outcome | Beta | P Value | FDR | 95% Confidence Interval |
| --- | --- | --- | --- | --- | --- |
| CN | Tau Tangle Pathology | -1.956 | 2.37E-07 | 7.54E-05 | -2.69 - -1.22 |
| CN | Neuritic Plaque Pathology | -0.829 | 4.08E-08 | 1.88E-05 | -1.12 - -0.54 |
| CN | Neurofibrillary Tangle Pathology | -0.584 | 8.29E-07 | 0.000289 | -0.81 - -0.35 |
| CN | Amyloid Pathology | -1.734 | 1.96E-07 | 0.000259 | -2.38 - -1.09 |
| DLPFC | Tau Tangle Pathology | -0.711 | 0.00125 | 0.0062 | -1.14 - -0.28 |
| DLPFC | Neuritic Plaque Pathology | -0.306 | 0.000479 | 0.00442 | -0.48 - -0.13 |
| DLPFC | Neurofibrillary Tangle Pathology | -0.176 | 0.00966 | 0.0398 | -0.31 - -0.04 |
| DLPFC | Amyloid Pathology | -0.293 | 0.124 | 0.238 | -0.67 - 0.08 |
| PCC | Tau Tangle Pathology | -1.509 | 0.000384 | 0.0136 | -2.34 - -0.68 |
| PCC | Neuritic Plaque Pathology | -0.757 | 2.04E-05 | 0.00209 | -1.1 - -0.41 |
| PCC | Neurofibrillary Tangle Pathology | -0.437 | 0.00108 | 0.0852 | -0.7 - -0.18 |
| PCC | Amyloid Pathology | -1.387 | 0.000274 | 0.0255 | -2.13 - -0.65 |

To profile the cell-type specific effect of *GSR* on human AD endophenotypes, we also performed analysis of ROSMAP snRNAseq (Synapse accession number: syn31512863). Differential expression patterns between individuals diagnosed with AD and normal cognition controls showed significant differences in *GSR* abundance in astrocytes (FDR = 0.04) but not in the remaining cell types including excitatory neurons, inhibitory neurons, oligodendrocytes, oligodendrocyte progenitor cells (OPCs), microglia, and endothelial cells (all FDR > 0.05). As the astrocyte association is modest, the snRNAseq results do not strongly support a specific cell type as the specific location for *GSR* expression-related effects on AD-related phenotypes.

Additional tests of predicted gene expression and tissue associations using PrediXcan models^33^ built with GTEx (v8 release, build 38) dataset (dbGaP accession number phs000424.GTEx.v8.p2 on 11/13/2019) did not show statistical significant results (all FDR > 0.05) (**Supplementary Table S3**).

## Discussion

We calculated a heritable quantitative measure of AD resilience, demonstrated its heritability, and mapped that trait in part to a region on chr8. We demonstrated that this locus was not significant for CFM in either Ntg or 5XFAD mice, identified a cis-pQTL for GR that appears to cleanly explain differences in resilience, and implicated protein networks related to cell and synaptic structure as well as protein folding and synapse organization that appear to manifest differentially in resilient animals. Overall, these data support a potential role for differential oxidative stress to confer resilience to AD pathology. We posit that the mechanism of differential oxidative stress tolerance is neuronal; it appears likely that neurons bearing the DBA/2J allele of GR have higher protein levels of the mouse *Gsr* gene and presumably greater reducing power, which allows them to tolerate greater oxidative stress than neurons with solely the C57BL/6J allele of GR. We speculate that the processes downstream of differential oxidative tolerance allow for neurons to remain intact against a variety of insults including those from amyloid plaque and tau tangle load.

The current dataset adds to a rich and mature literature on redox balance as a potential mechanism of AD that suggests redox balance as a potential mechanism for resilience. This literature primarily derives from biochemistry. The AD brain undergoes a variety of oxidative insults^34^, and glutathione redox balance has long been suggested as one potential mechanism for oxidative stress in neurodegenerative diseases^35^. Glutathione redox state itself has been the focus of AD in recent decades^36^, though difficulty in assessing redox state in postmortem samples has likely prevented direct assessment of the ratio of GSSG to GSH in AD brains. A recent quantitative magnetic spectroscopy study identified a decreased abundance of GSH in AD patient hippocampus relative to healthy subjects *in vivo*^37^. Thus, the literature supports a biochemical role for glutathione balance as a potential mechanism or consequence of AD pathology, which suggests that alteration of glutathione redox balance may confer resilience.

Though the biochemical evidence supports redox as a potential AD mechanism or consequence, human genetics has not strongly supported this hypothesis. Candidate gene studies have suggested glutathione metabolism as an AD mechanism^38,39^. However, to our knowledge, this is the first time that a genetics study has directly supported glutathione redox state as a genome-wide significant AD finding. Neither the most exhaustive recent human GWAS on AD^40^ nor the most exhaustive recent human GWAS on AD resilience^41^ identified redox or mitochondrial metabolism as a potential genetic mechanism in humans. We note that by using a recombinant inbred mouse population, we boost mappable heritability through repeated sampling^14^. This property enhances our ability to ascertain lower heritability resilience genetics, a property that is not available in any human population. Further, we note several possibilities for the lack of human genetics findings supporting glutathione redox state. First, the specific mechanism we observed may not be part of the common genetic variation in widely studied human population and is thus not resolvable by GWAS. Second, it is possible that common variation does exist for this specific mechanism, but that low heritability means that even large GWAS may be underpowered for these loci. Third, it is possible that a search for rare human variants in *GSR* that may confer resilience to AD, a strategy that has recently identified a variant of interest in *REELIN*^7^. Finally, mice afford us better environmental and nutritional control, which reduces the environmental noise relative to human experiments.

In support of our hypothesis about rare genetic variation, mutations in the *GSR* gene are rare in human populations. Two polymorphisms in the gene region have been associated with a decrease in human GR activity: a missense variant, rs8190955_A (p = 4 x 10^-86^) ^42^ and an intron variant, rs4733505_A (p = 1 x 10^-26^) ^43^. We note that while the highly penetrant missense variant rs8190955 is rare with a frequency of 0.87% in the human population as a whole, and 0.25% in the European population, these mutations are more frequent in those of other ancestries that are now becoming better represented in GWAS (African: > 7.4% and Asian: > 4.2%). As the population and European population frequencies are below the minor allele frequency threshold used in many GWAS analysis pipelines, it is possible that ascertainment issues have prevented GWAS from detecting an association that is observed in mice, further reiterating the benefits of studying both diverse mouse and human populations in research.

Human functional genomics findings do support a role for glutathione S reductase. We found that the human *GSR* gene expression and cognition findings in the ROSMAP study presented here matches the direction of the mouse results. Increased *Gsr* expression is correlated with higher cognitive function and lower AD pathology, offering evidence of translation to humans. As in mice, the human GR protein plays a role in restoring glutathione to its reduced state (GSH).

Reduced glutathione acts an antioxidant with demonstrated neuroprotective effects in humans^44^. Additionally, depleted mitochondrial GSH (mGSH) has led to Aβ-induced neuroinflammation and neurotoxicity, effects reversed by mGSH replenishment^45^. Similar observations for mGSH were observed in mouse models^46^.

This work suggests several potential novel pharmaceutical targets that might act as mimetics of this resilience mechanism. For example, GR itself is the target of several compounds developed as potential antimalarials^47^, suggesting that such compounds may lead to the creation of a new generation of small molecules targeting GR. Further, evidence that oxidative damage is a part of AD provides support for the potential of blood-brain barrier permeable mitochondrial antioxidants as potential novel AD therapeutics, a hypothesis that has some history^48,49^.

Finally, the effects we observed here are only the first and most notable effects that might be derived from these data. Not only is the metric we used to measure resilience novel, the proteomics dataset itself is a novel resource that can be deeply mined. We believe these data will act as a rich resource aiding further study, assisting in the interpretation and elucidation of various aspects of normal aging hippocampus as well as AD pathology.

## Materials and Methods

### Animals

All mice used in the study were housed in the University of Tennessee Health Science Center or at The Jackson Laboratory under 12:12 light/dark cycle with food and water ad libitum. F1 Ntg-BXD and AD-BXDs used in this paper were generated as previously described^9^. Briefly, female congenic C57BL/6J mice hemizygous to the 5XFAD transgenes were crossed to males from the recombinant inbred BXD mouse genetic reference panel. The resulting F1 population inherits a single BXD allele and either a single C57BL/6J with the 5XFAD transgenes (AD-BXDs) or C57BL/6J allele without the 5XFAD transgenes (Ntg-BXD). Confirmation of 5XFAD transgene carrier status was performed using genotyping either at The Jackson Laboratory Transgenic Genotyping Services or Transnetyx (TN, USA). All procedures were carried out in accordance with the standards of the Association for the Assessment and Accreditation of Laboratory Animal Care (AAALAC) and the National Institutes of Health (NIH) Guide for the Care and Use of Laboratory Animals and were approved by the respective Institutional Animal Care and Use Committee (IACUC).

### Behavioral Data and Tissue Collection

Prior to tissue collection, animals underwent contextual fear conditioning and Y maze behavioral testing as described previously ^9^. Animals in this study were aged to either 6 months or 14 months. Y maze data were collected at both 6 month and 14 month time points while contextual fear conditioning was performed just prior to euthanasia. Hippocampal and frontal cortex samples were harvested from 6 month and 14 month AD-BXDs and Ntg-BXDs and were snap frozen immediately following fear conditioning and were shipped to the University of Washington for DIA proteomics.

### DIA Proteomics

The mouse hippocampus tissue was randomly balanced based on condition group ratios (genotype, sex, age and BXD strain) into batches of 14 individual hippocampal samples and 2 references. One reference was the same pool of a balanced batch to create a hippocampus specific reference that could be used as a common reference or a single-point calibrant in every batch to not only monitor the performance of the experiment workflow but also calibrate variability across different laboratories or experiments^50^. To evaluate the run order and batch effects within the study, a second reference, which is a mixture of C57BL/6J (B6/B6) and DBA/2J (B6/D2) mouse midbrain and cerebellum tissues from all condition types, was used as a batch reference to allow for a more accurate comparison of between brain region proteomic profiles. The hippocampal tissue was processed as previously described^51^. In summary, the tissue was briefly probe sonicated in a SDS lysis buffer and then a small volume of lysate was high pressure homogenized. In order to monitor digestion, a process control of yeast enolase was added to the protein homogenate which was then reduced, alkylated and trypsin digested on a S-trap column (Protifi). Eluates of hydrophilic and hydrophobic peptides were pooled and speed vacuumed. One μg of each digested sample, 8 ng of yeast enolase and 150 femtomole of Pierce Retention Time Calibrant (PRTC) were loaded onto a Thermo EASY nano-flow UHPLC coupled with a Thermo Orbitrap Fusion Lumos Mass Spectrometer. The yeast enolase is used as a protein process control and the PRTC is used as a peptide process control within the sample. System suitability (QC) injections of 150 fmol of PRTC and BSA were also used independently, to assess quality before and during analysis. We analyzed four of these system suitability runs prior to any sample analysis and then after every six to eight sample runs another system suitability run is analyzed. Six gas phase fractionated LC-MS/MS DIA runs of an overlapping “narrow” 4 m/z isolation window of a mass range of 100 m/z (400-500 m/z, 500-600 m/z, 600-700 m/z, 700-800 m/z, 800-900 m/z, 900-1000 m/z) for each of the six runs from a pool of all the samples from each batch are collected to be used as the on-column chromatogram library. Each individual digested sample was also analyzed with a LC-MS/MS DIA run with a mass range of 400-1000 m/z and an overlapping “wide” 8 m/z isolation window. The quality control data was analyzed using Skyline and AutoQC^52,53^.

### Signal Processing

Thermo instrument RAW files were converted to mzML format using Proteowizard (version 3.0.20064) using vendor peak picking and demultiplexing with the settings of “overlap_only” and Mass Error = 10.0 ppm^54^. EncyclopeDIA (version 2.12.30) was used to search and combine the on column chromatogram libraries of all runs from all 31 batches using the data from the six gas phase fractionated “narrow window” DIA runs of a pool from each batch^22^. These narrow windows were analyzed using EncyclopeDIA with the default settings (10 ppm tolerances, trypsin digestion, HCD b- and y-ions) of a Prosit predicted spectra library based on the Uniprot mouse canonical FASTA (April 2020) ^55^. The results from this analysis were saved as a “Chromatogram Library’’ in EncyclopeDIA’s eLib format where the predicted intensities and iRT of the Prosit library were replaced with the empirically measured intensities and RT from the gas phase fractionated LC-MS/MS data^56^. The “wide window” DIA runs were analyzed using EncyclopeDIA (version 2.12.30) requiring a minimum of 3 quantitative ions and filtering peptides with q-value ≤ 0.01 using Percolator (version 3.01). After analyzing each file individually, EncyclopeDIA was used to generate a “Quant Report’’ which stores all of the detected peptides, integration boundaries, quantitative transitions, and statistical metrics from all runs in an eLib format. The Quant Report eLib library is imported into Skyline (version 22.2.0.351) with the mouse uniprot FASTA as the background proteome to map peptides to proteins, perform peak integration, manual evaluation, and report generation. A csv file of peptide level total area fragments (TAFs) for each replicate was exported from Skyline using the custom reporting capabilities of the document grid^52^.

### Post-processing

The quantitative peptide data exported from Skyline (level 2 data) was post-processed to minimize residual technical noise. The data was log2 transformed and median normalized to scale the intensities of the sample to the same median value. Using the batch factor as a predictor variable and abundance as its response, a simple linear regression model is fit to model the confounding due to the batch effect bias. The residuals from this model are assumed to represent the peptide or protein abundance without the effect of the batch covariate. Principal Component Analysis (PCA) is used to evaluate the confounding bias by projecting the data in its reduced dimensions onto the first three principal components. The normalized and batch adjusted peptide abundance is provided as level 3A data. Peptides are next constructed into an indistinguishable protein group and are summed to estimate the abundance of the peptide groups and proteins that match the same set of peptide groups merged into a single node in the graph^57,58^. The normalized and batch adjusted protein abundance are provided as level 3B data.

### Resilience Quantitative Trait

To compute resilience, we utilized the property of trait correlation within the BXD panel. This property allowed us to compare the cognitive performance (contextual fear memory, CFM) of AD-BXD mice bearing a 5XFAD transgene with corresponding strain of Ntg mice bearing no AD risk locus. This technique allows us to assess cognitive performance of an animal with an AD risk transgene relative to the same strain without the AD risk transgene.

Briefly, we took the average CFM score for each Ntg strain. We then regressed the CFM score for 5XFAD AD-BXD mice of the same genotype against these averages using weighted regression. Weights used were 1/*n* where *n* is the sample size within each 5XFAD AD-BXD strain. This technique fits a line identical in slope and intercept to a regression of strain means, but allows us to compute individual residuals for each 5XFAD AD-BXD mouse in the dataset. We then took the standardized residuals, defined as the Z scores of the residuals, and defined these scores as the quantitative resilience trait.

### Heritability

Heritability was calculated using the sum of squares of an ANOVA model with BXD strain as the independent variable. This calculation was performed for any strain for which a measure had a sample size of *n* ≥ 2. This heritability calculation is as follows:

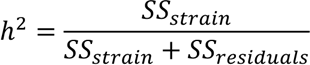

For recombinant inbred panels, measured heritability of a phenotype may be boosted by taking the strain mean heritability value^14^, which accounts for multiple measurements in the same strain. Strain mean heritability was calculated from the above heritability value using the following calculation:

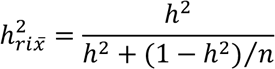

where *n* is the average sample size per strain in the experiment.

### Resilience Mapping

Mapping of the resilience quantitative trait was performed using a univariate linear mixed model as implemented in GEMMA v0.98.6^59^. Strain means of the resilience quantitative trait were first computed and rank-Z transformed for 14-month aged 5XFAD AD-BXD mice. We used the 2017 BXD genotypes accessed from https://gn1.genenetwork.org/dbdoc/BXDGeno.html. Coordinate space was lifted over to the GRCm39/mm39 genome space using the `liftOver` R package with the appropriate chain file from UCSC (`mm10ToMm39.over.chain`). One marker (`14P_no_data`) did not lift over and was manually assigned a GRCm39 coordinate of chr3:3.000000 Mb. We computed separate kinship matrices for each chromosome using GEMMA’s inbuilt leave-one-chromosome-out (LOCO) kinship estimation algorithm.

The final mapping model in GEMMA utilized the rank-Z transformed data, the GRCm39 lifted over genotype data, and the LOCO kinship estimates. LOD scores were calculated from p-values by taking −*log*_10_(*p*). Mapping results were additionally confirmed using R/qtl2 v0.30^60^ in parallel and with parallel settings. A LOD score threshold of 0.01 was calculated from 1,000 permutations of the data using the `scan1perm()` function. Peaks were identified using the function `find_peaks()` on the R/qtl2 confirmatory mapping results with a 99% Bayes confidence interval.

### Protein Abundance Analysis

To identify the presumptive protein, we performed a differential expression analysis of protein expression values as a consequence of the peak resilience marker on chr8 (rs33539160).

Briefly, we modeled expression as a consequence of the additive covariate of sex and of genotype at rs33539160. Differential analysis was performed on individual 14-month aged 5XFAD AD-BXD samples using limma v3.54.2^61^. Results were false discovery rate adjusted using q values^62^ as implemented in the qvalue package v2.30.2. Mediation analysis was performed using the `mediate` function in the mediation R package v 4.5.0 with 30,000 simulations using the quasi-Bayesian approximation method.

### pQTL Analysis

pQTL analysis utilized the same version of GEMMA and the same genotype files as in the resilience mapping analysis. However, due to the sparsity of proteomics data, an additional criterion was needed: pQTL were only calculated for proteins with at least 10 strains for which measurements existed. Peaks with LOD scores greater than 4 and a between-peak LOD score drops of 1.5 were computed using the `qtl2:::.find_peaks()` function from R/qtl2 with the appropriate adjustment made for differences in indexing between the base C++ function and R. Peaks were called as cis-pQTLs if their protein was encoded by a gene on the same chromosome as a peak and if the peak was within 25 Mb of that gene.

To identify probable trans relationships with GR, we found any trans-pQTL that mapped within 15 Mb of the peak marker for GR. We used the UniProt protein IDs for these proteins as their names.

### Paraclique Analysis and GO Enrichment

We used the Paraclique algorithm^29^ as implemented in the cliqueR R package v1.0. Briefly, Paraclique identifies network cliques, subnetworks in which all nodes are adjacent to one another. It then expands this subnetwork using a glom factor that governs the degree of additional nodes added to the cluster. The result is a set of coexpression paracliques, densely clustered coexpression modules that are ordered in ascending order by how many nodes they have. Unlike other methods, there is no requirement for mutually exclusive clusters in paracliques.

To address the sparsity of the proteomics data, we used the `ppca` method implemented in the pcaMethods package v1.90.0^63^ to impute missing expression values. We used the `bicor` function from the WGCNA package v1.72-1^64^ to compute correlation coefficients and filtered coexpression relationships to those with a bicor ≥ 0.40, equivalent to a signed analysis in WGCNA. Paracliques were computed with a minimum clique size of 5, a minimum paraclique size of 10, and a cliqueR glom factor of 0.2. Eigengenes were computed for each paraclique by computing the first principal component of protein expression values from each cluster. We analyzed the relationship of the clusters to resilience by linear regression with the quantitative resilience scores. The p-values from the regression were corrected for false discovery rate using the qvalue package v2.30.2 and relationships of q < 0.10 were called significant.

Visualization utilized a method inspired by REVIGO^65^ in which semantic similarity of GO BP terms was used as the input for a distance matrix, which was then subjected to multidimensional scaling. Briefly, we used the PANTHER website (https://pantherdb.org/) to perform GO enrichment analysis on UniProt IDs using the PANTHER GO-Slim Biological Process (GO BP) dataset. Significance values for GO BP categories were corrected using the Bonferroni method. Semantic similarity between significant GO BP categories was computed using the GOSemSim v2.24.0 R package^66^ using the Wang algorithm^67^. Similarity measures were used to build a dissimilarity matrix for non-metric multi-dimensional scaling using the `isoMDS` function from the MASS R package v7.3-57 for visualization of GO clusters. Clusters were called using the `cutreeDynamicTree` function from the dynamicTreeCut R package v1.63-1.

### Human *GSR* Transcript Abuncance Association with AD Pathology Using ROSMAP Bulk RNAseq Data

All ROSMAP participants enrolled without known dementia and agreed to detailed clinical evaluation and brain donation at death. Both studies were approved by an Institutional Review Board of Rush University Medical Center. Each participant signed an informed consent, Anatomic Gift Act, and repository consent to allow their data to be repurposed.

Processed bulk RNAseq data^32,68,69^ from 3 tissues – dorsolateral prefrontal cortex (DLPFC), posterior cingulate cortex (PCC), and the head of the caudate nucleus (CN) – were used to build multiple linear regression models (for cross-sectional outcomes) and linear mixed-effect models (for longitudinal outcomes) for testing *GSR* gene expression associations with AD pathology, including β-amyloid load by immunohistochemistry (IHC), neurofibrillary tangles by IHC, as well as longitudinal measures cognitive function. Global cognitive score was the main variable used to represent overall cognitive function. This score which was computed by converting raw scores from 19 cognitive tests to Z scores and then averaged. For the cross-sectional cognitive function related analysis, the global cognitive score at last visit before death was used. For the longitudinal model, a cognitive trajectory was quantified from a mixed effects regression model with both the intercept and the slope entered as fixed and random effects and global cognition set as the outcome. The cognitive trajectory was extracted as the estimated person-specific annual rate of change in global cognition. Covariates included in these models were age at death, sex, postmortem interval (PMI), and interval between last visit and death for all cross-sectional outcomes. In longitudinal models, time was modeled as the interval between a visit and the last visit, calculated in years.

### ROSMAP Single Nucleus RNAseq (snRNAseq)

snRNAseq data of DLPFC brain specimens derived from 424 ROSMAP participants (Synapse accession number: syn31512863) were generated and processed at Columbia University Medical Center^70^. There are in total 8 cell types used in our analysis: cux2+/cux2- excitatory neurons, inhibitory neurons, astrocytes, microglia, oligodendrocytes, oligodendrocyte precursor cells (OPCs), and endothelial cells. Genes with expression in a minimum of 10% of all cells were used for modeling. Cells were removed if they counted more than 20,000 or less than 200 total RNA UMIs, or had more than 5% mitochondrially mapped reads. The gene count matrix used was the UMI count data from RNA assay normalized and scaled by “sctransform” R package (https://github.com/satijalab/sctransform).

All association analysis was carried out in cell-type specific manner using negative binomial lognormal mixed models implemented in the NEBULA R package (v1.2.0) ^71^. Differential gene expression was profiled between participants with normal cognition (n= 142) at last clinic visit and AD dementia groups (n = 157), covarying for age at death, sex, and PMI. The same covariates were used for estimating the association with AD outcomes, including β-amyloid (IHC staining), neurofibrillary tangles (IHC staining), both cross-sectional and longitudinal cognitive function. Please refer to bulk RNAseq method for the definition of cognitive function.

All statistical analyses were carried out using R (v4.2.1). Significance was set *a priori* to α = 0.05. P values were corrected for multiple comparisons using the Benjamini-Hochberg false discovery rate procedure ^72^ across all genes and outcomes tested.

## Acknowledgments

This work was supported by NIH R01 AG057914 and R01 AG075818 to CCK, P30 AG013280 and U19 AG065156 to MJM, and NIH R01 AG061518 and R01 AG074012 to TJH. ROSMAP is supported by P30 AG010161, P30 AG072975, R01 AG015819 and R01 AG017917 to DAB as well as U01 AG046152 and U01 AG061356 to DAB and PLDJ.

## Data Availability

The Skyline documents, raw files for quality control and DIA data are available at Panorama Public. ProteomeXchange ID: PXD045425. DOI: https://doi.org/10.6069/qsmg-5m62. Access URL: https://panoramaweb.org/AD-BXD-mouse-hipp-proteomics.url^73^.

DIA data is available in 5 different categories based on the level of post-processing. Level 0 represents the raw data in two different formats - the raw format is directly from the Thermo mass spectrometer and the mzML format is the demultiplexed version of the raw data (Proteowizard version 3.0.20064). Level 1 describes the zipped Skyline document. Level 2 is a csv file of the Skyline output with the integrated peak area (total area fragments) for each peptide (row) in each replicate (column). Level 3A is a csv file of the normalized peptide abundance across all batches. Level 3B is a csv file of the normalized protein abundance across all batches.

Quality control Skyline documents, peptide QC plots and instrument raw files for system suitability runs are provided for all 31 batches. The Skyline documents and peptide QC plots for enolase and PRTC process controls are provided by column batch. The instrument raw files for process controls are the same as DIA sample raw files by column batch.

ROSMAP data are available online on the Accelerating Medicines Partnership – Alzheimer’s disease (AMP-AD) Knowledge Portal (syn3219045 and syn31512863) or through the Rush Alzheimer’s Disease Center Resource Sharing Hub (https://www.radc.rush.edu/).

