## Supplementary Table S3 for "Hippocampus Glutathione S Reductase Potentially Confers Genetic Resilience to Cognitive Decline in the AD-BXD Mouse Population": supplementary_table_s3.html

GSR gene lookup


Code 

- Show All Code
- Hide All Code
- Download Rmd

### GSR gene lookup

###### Yiyang Wu - Hohman Lab @VUMC

###### Created: Sep 2023 | Last Modified: September 12,2023


### 1 bulkRNAseq from ROSMAP

#### 1.1 Background

The analysis shows results from ROS/MAP longitudinal study and
autopsy. Data sets include the caudate nucleus (CN), dorsolateral
prefrontal cortex (DLPFC), and posterior cingulate cortex (PCC). The
analysis question is, do the gene expressions have an association with
cognition decline and AD-pathology. To evaluate the data, a multiple
linear regression model (cross-sectional and AD-related pathology) as
well as a linear mixed-effects model (longitudinal) was used.

**Model Overview**

**Outcome**

Global Cognition Before Last Visit

Longitudinal Global Cognition

Autopsy Path (Amyloid, Neuritic Plaques (NP), Tangles,
Neurofibrillary Tangles (NFT))

**Covariates**

Age at Death, Sex, Post-Mortem Interval (PMI), Interval Death,
Latency to Death

#### 1.2 Demographic Information

Below is the demographic information for genes in ROS/MAP.


```
# Use make_descriptive_table to make the table descriptives for each gene and tissue
make_descriptive_table(last_visit, ENSG_code, gene_name)
```


| ENSG\_code | Tissue | CN | DLPFC | PCC |
| --- | --- | --- | --- | --- |
| GSR | | | | |
| --- | --- | --- | --- | --- |
| ENSG00000104687 | Education | 16.32±3.62 | 16.39±3.56 | 16.4±3.57 |
| ENSG00000104687 | % Male | 33% | 32% | 37% |
| ENSG00000104687 | % Normal cognition | 35% | 35% | 38% |
| ENSG00000104687 | Age at death | 89.22±6.49 | 89.38±6.68 | 89.31±6.54 |
| ENSG00000104687 | Global cognition | -0.76±1.07 | -0.8±1.07 | -0.68±1.03 |
| ENSG00000104687 | N | 701 | 944 | 524 |
| ENSG00000104687 | PMI | 7.62±4.42 | 7.57±4.34 | 7.05±4.04 |

#### 1.3 Histograms


```
# Use make_histograms function to create a histogram for each gene per tissue
make_histograms(last_visit, ENSG_code, gene_name)
```

#### 1.4 Summary Result of the Association Analysis (cross-sectional & longitudinal)

##### 1.4.1 Cognition Outcome


```
# Grabs files in the folder
list_of_files <- list.files(paste0(path, "input/RegressionModels/cognition/combine/")
  )

df <- lapply(1:length(list_of_files), function(i) {
  df <-
    data.table::fread(
      paste0(
        path, "input/RegressionModels/cognition/combine/",
        list_of_files[i]
      )
    )
  df %>% dplyr::filter(Ensembl_gene %in%c(ENSG_code))
})

dfs <- do.call("rbind", df) # combine data frames

# calculate an FDR correction for the multiple models and tissues
all_results_main_effect <- dfs %>%
  dplyr::select("Ensembl_gene", "Tissue", "Outcome", "Beta", "StdError",
                "DF", "Pvalue", "P.fdr")

 # making data frame with ENSG code and gene name
  gene_info <- as.data.frame(ENSG_code, gene_name)
  
  # using tibble row name to column
  gene_info <- rownames_to_column(gene_info, var = "Gene")
  
  # renaming gene_info first column to Ensembl_gene
  colnames(gene_info)[2] <- "Ensembl_gene"
  
  # left joining gene_info with the combined_description data
  final_combined_data <-
    left_join(all_results_main_effect, gene_info, by = "Ensembl_gene")
  
  # remove the ENSG code column
  final_combined_genes <- final_combined_data %>%
    dplyr::select(-Ensembl_gene) %>% 
    dplyr::select(Gene, everything())

all_results_main_effect <-
  format(final_combined_genes, digits = 3) # change decimals

# create datatable with 10 page length
DT::datatable(all_results_main_effect,
              rownames = FALSE,
              filter = 'top',
              extensions = 'Buttons',
              options = list(pageLength = 10, 
                             dom = 'Bfrtip',
    buttons = c('copy', 'csv', 'print'))) %>%
  DT::formatStyle(c("P.fdr"),
                  color = DT::styleInterval(0.05, c("red", "black")))
```

##### 1.4.2 Pathology Outcome


```
# Grabs files in the folder
list_of_files <-
  list.files(
    paste0(path, "input/RegressionModels/pathology/combine/")
  )

df <- lapply(1:length(list_of_files), function(i) {
  df <-
    data.table::fread(
      paste0(path, "input/RegressionModels/pathology/combine/",
        list_of_files[i]
      )
    )
  df %>% dplyr::filter(Ensembl_gene %in%c(ENSG_code))
})

dfs <- do.call("rbind", df) # combine data frames

# calculate an FDR correction for the multiple models and tissues
all_results_pathology <- dfs %>%
  dplyr::select("Ensembl_gene", "Tissue", "Outcome", "Beta", "StdError",
                "DF", "Pvalue", "P.fdr")

 # making data frame with ENSG code and gene name
  gene_info <- as.data.frame(ENSG_code, gene_name)
  
  # using tibble row name to column
  gene_info <- rownames_to_column(gene_info, var = "Gene")
  
  # renaming gene_info first column to Ensembl_gene
  colnames(gene_info)[2] <- "Ensembl_gene"
  
  # left joining gene_info with the combined_description data
  final_combined_data <-
    left_join(all_results_pathology, gene_info, by = "Ensembl_gene")
  
  # remove the ENSG code column
  final_combined_genes <- final_combined_data %>%
    dplyr::select(-Ensembl_gene) %>% 
    dplyr::select(Gene, everything())

all_results_pathology <-
  format(final_combined_genes, digits = 3) # change decimals

# create datatable with 10 page length
DT::datatable(all_results_pathology,
              rownames = FALSE,
              filter = 'top',
              extensions = 'Buttons',
              options = list(pageLength = 10,
                             dom = 'Bfrtip',
    buttons = c('copy', 'csv', 'print'))) %>%
  DT::formatStyle(c("P.fdr"),
                  color = DT::styleInterval(0.05, c("red", "black")))
```

#### 1.5 Plots

##### 1.5.1 Volcano Plots

Below are the volcano plots for each tissue and each outcome of
interest. The first layer of the volcano plot is black, representing all
the genes. The second layer is green, representing any genes whose
p-values reached significance level of less than 0.05. The third layer
is the red dot, displaying where the gene of interest is located on the
plot.

The AdjustedR2 value was derived from subtracting the marginal R2
value from the adjustedR2 value. If the beta value was less than zero,
the resulting R2 was multipled by negative one.


```
makeVolcanoPlotsAmyloid(
  paste0(
    path,
    "input/RegressionModels/ByTissue/combine/all_tissue_results.csv"
  ),
  ENSG_code,
  gene_name,
  gene_names_codes
)
```


```
|--------------------------------------------------|
|==================================================|
```


```
makeVolcanoPlotsCrossSectional(
  paste0(
    path,
    "input/RegressionModels/ByTissue/combine/all_tissue_results.csv"
  ),
  ENSG_code,
  gene_name,
  gene_names_codes
)
```


```
makeVolcanoPlotsLongitudinal(
  paste0(
    path,
    "input/RegressionModels/ByTissue/combine/all_tissue_results.csv"
  ),
  ENSG_code,
  gene_name,
  gene_names_codes
)
```


```
makeVolcanoPlotsNFT(
  paste0(
    path,
    "input/RegressionModels/ByTissue/combine/all_tissue_results.csv"
  ),
  ENSG_code,
  gene_name,
  gene_names_codes
)
```


```
makeVolcanoPlotsNP(
  paste0(
    path,
    "input/RegressionModels/ByTissue/combine/all_tissue_results.csv"
  ),
  ENSG_code,
  gene_name,
  gene_names_codes
)
```


```
makeVolcanoPlotsTauTangle(
  paste0(
    path,
    "input/RegressionModels/ByTissue/combine/all_tissue_results.csv"
  ),
  ENSG_code,
  gene_name,
  gene_names_codes
)
```

### 2 snRNAseq ROSMAP\_syn2580853

#### 2.1 Background of the Study

This project utilized recently published single-cell transcriptomes
derived from dorsolateral prefrontal cortex tissues of post-qc 424
donors collected at Rush Alzheimer’s Disease Center. Processed single
cell RNA sequencing data were acquired from AD Knowledge Portal
(syn2580853). Mean age at death was 89 years, 68% were females, and 37%
were AD cases.

#### 2.2 DGE Analysis Between Normal Cognition Control and AD Individuals

##### 2.2.1 Method

299 individuals with normal cognition control (n= 142, cogdx = 1 at
initial clinic visit) or AD dementia (n = 157, cogdx = 4 or 5 at initial
clinic visit) were analyzed for differential gene expression in all 8
cell types. Negative binomial lognormal mixed models implemented in the
NEBULA R package (v1.2.0) were used for this analysis, with the model
“count matrix ~ sex + age\_death + pmi + clinical group”. Genes with
expressions in minimum 10% of all cells were used for the model. Cells
were removed if they counted more than 20,000 or less than 200 total RNA
UMIs, or had more than 5% mitochondrially mapped reads. The gene count
matrix input of the model was the UMI count data from RNA assay
normalized and scaled by “sctransform” R package (https://github.com/satijalab/sctransform). NEBULA-HL
method is used for the modeling. The p value calculated from the models
was FDR adjusted using R function “p.adjust” using “BH” as the
method.

Abbreviations of cell types: ex = excitatory neurons, inhib =
inhibitory neurons, oligo = oligodendrocytes, opc = oligodendrocyte
progenitor cells, micro = microglia, endo = endothelial cells, ast =
astrocytes

##### 2.2.2 Summary Result


```
DE <- readRDS(paste0(snRNA_input, "DE_Nebula_01312023.rds"))
# DE <- data.frame(DE) %>%
#   dplyr::mutate(logFC_AD_over_Control = logFC_Control_over_AD*(-1)) %>%
#   dplyr::select(c(1,2,7,4:6)) %>%
#   setNames(c('Gene', 'Cell_type', 'logFC_AD_over_Control', 'SE', 'P', 'FDR'))
# saveRDS(DE, paste0(snRNA_input, "DE_Nebula_AD_over_ctrl_08082023.rds"))
```


```
DE_table <- DE %>%
  dplyr::filter(gene %in% gene_name) %>%
  dplyr::arrange(gene)

DT::datatable(DE_table, 
              rownames = FALSE,
              filter = 'top',
              extensions = 'Buttons',
              options = list(pageLength = 10,
                             dom = 'Bfrtip',
    buttons = c('copy', 'csv', 'print'))) %>% 
      DT::formatRound(3:6, digits = 3) %>% 
      DT::formatStyle(c("FDR"),
                  color = DT::styleInterval(0.05, c("red", "black")))
```


###### 2.2.2.0.1 plot


```
makeVolcanoPlot_nblmm(
  paste0(snRNA_input, "DE_Nebula_01312023.rds"),
  gene_name
)
```

#### 2.3 Association Analysis between Gene Expression and 4 Quantitative Traits

##### 2.3.1 Method

To estimate the association between gene expression and amyloid and
tangles outcomes, while accounting for sex, age of death, and postmortem
interval (pmi), the following model formulation were used respectively;
“count matrix ~ amyloid + sex + age of death + pmi”, count matrix ~
tangles + sex + age of death + pmi”. To estimate the association between
gene expression and cross-sectional last visit cognition global scores,
while accounting for interval between last visit and death, sex, age of
death, and pmi, the following model formulation was used; “count matrix
~ global cognitive score at last visit + sex + age of death + pmi +
interval between last visit and death”, with interval modeled as time in
years. To estimate the association between gene expression and
longitudinal cognition decline, while accounting for interval between
last visit and death, sex, age of death, and pmi, the following model
formulation was used; “count matrix ~ cognition slope + sex + age of
death + pmi + interval between last visit and death”.

##### 2.3.2 Summary Result

###### 2.3.2.1 Cross-sectional Cognition


```
cross_cog <- readRDS(paste0(snRNA_input, "last_cognition_Nebula_01312023.rds"))
cross_cog <- data.frame(cross_cog)
```


```
cross_cog_table <- cross_cog %>%
  dplyr::filter(gene %in% gene_name) %>%
  dplyr::arrange(gene)

DT::datatable(cross_cog_table, 
              rownames = FALSE,
              filter = 'top',
              extensions = 'Buttons',
              options = list(pageLength = 10,
                             dom = 'Bfrtip',
    buttons = c('copy', 'csv', 'print'))) %>% 
      DT::formatRound(3:6, digits = 3) %>%
      DT::formatStyle(c("P_FDR"),
                  color = DT::styleInterval(0.05, c("red", "black")))
```

###### 2.3.2.2 Longitudinal Cognition


```
long_cog <- readRDS(paste0(snRNA_input, "long_cognition_Nebula_01312023.rds"))
long_cog <- data.frame(long_cog)
```


```
long_cog_table <- long_cog %>%
  dplyr::filter(gene %in% gene_name) %>%
  dplyr::arrange(gene)

DT::datatable(long_cog_table, 
              rownames = FALSE,
              filter = 'top',
              extensions = 'Buttons',
              options = list(pageLength = 10,
                             dom = 'Bfrtip',
    buttons = c('copy', 'csv', 'print'))) %>% 
      DT::formatRound(3:6, digits = 3) %>%
      DT::formatStyle(c("P_FDR"),
                  color = DT::styleInterval(0.05, c("red", "black")))
```

###### 2.3.2.3 Immunohistochemical Amyloid


```
amy <- readRDS(paste0(snRNA_input, "amyloid_Nebula_01312023.rds"))
amy <- data.frame(amy)
```


```
amy_table <- amy %>%
  dplyr::filter(gene %in% gene_name) %>%
  dplyr::arrange(gene)

DT::datatable(amy_table, 
              rownames = FALSE,
              filter = 'top',
              extensions = 'Buttons',
              options = list(pageLength = 10,
                             dom = 'Bfrtip',
    buttons = c('copy', 'csv', 'print'))) %>% 
      DT::formatRound(3:6, digits = 3) %>% 
      DT::formatStyle(c("P_FDR"),
                  color = DT::styleInterval(0.05, c("red", "black")))
```

###### 2.3.2.4 Immunohistochemical Tangles


```
tau <- readRDS(paste0(snRNA_input, "tangles_Nebula_01312023.rds"))
tau <- data.frame(tau)
```


```
tau_table <- tau %>%
  dplyr::filter(gene %in% gene_name) %>%
  dplyr::arrange(gene)

DT::datatable(tau_table, 
              rownames = FALSE,
              filter = 'top',
              extensions = 'Buttons',
              options = list(pageLength = 10,
                             dom = 'Bfrtip',
    buttons = c('copy', 'csv', 'print'))) %>% 
      DT::formatRound(3:6, digits = 3) %>%
      DT::formatStyle(c("P_FDR"),
                  color = DT::styleInterval(0.05, c("red", "black")))
```

### 3 PrediXcan

#### 3.1 Predicted Expression - Resilience

##### 3.1.1 Method

Gene expression association analysis was performed using gene
expression imputed by applying tissue specific (49 tissues with on an
average of 5455 genes examined) PrediXcan models built with GTEx (v8
release, build 38) dataset (dbGaP accession number phs000424.GTEx.v8.p2
on 11/13/2019), an elastic-net supervised machine learning model (α =
0.5, equal weights for lasso and ridge regression) following the method
described in Gamazon et al (Nature Genetics 2015) and applied to
genotypes. Geneotypes were prepared with the established GWAS\_QC
pipeline. Briefly raw genotypes with minor allele frequency
(MAF)<0.01 were excluded, indels were removed and all variants were
lifted to genome build 38. After restricting to Non-Hispanic White
individuals (WGS genotypes were not imputed), genotypes were imputed on
TOPMed -r2 panel on TOPMed imputations server (https://imputation.biodatacatalyst.nhlbi.nih.gov ).
Following imputation, variants were restricted to bi-allelic SNPs and
limited to those with an imputation R2>0.8 and MAF>0.01. RNA
expression values were normalized, following the GTEx consortium’s
pipeline (https://github.com/broadinstitute/gtexpipeline/blob/master/TOPMed\_RNAseq\_pipeline.md).
Briefly, across samples, values were normalized to the average empirical
distribution and then, within each gene, values were inverse quantile
normalized. Genes expressed in at least 10 samples at greater than 0.1
RPKM and greater than 5 reads were carried forward. Then, expression
values were corrected using a residual method for 60 PEER factors, sex,
genetic principal components, and sequencing platform. In summary, for
whole blood expression for 26,946 genes in 567 samples was leveraged for
building models of predicted expression.”

**Outcome**

Combined Cognitive Resilience (GLOBALRES)

Combined Cognitive Resilience Normal Cognition (GLOBALRES NC)

**Covariates**

Age at Visit

Gender

**Models**

GLOBALRES ~ age + gender + list of genes

GLOBALRES NC ~ age + gender + list of genes

##### 3.1.2 Tissue Information Table

#### 3.1.2.1 A4


```
A4_table <-
  readr::read_csv(
    paste0(path, "input/PrediXcanTissueTables/A4_predicted_expression_qc.csv")
  )

A4_table <- A4_table %>%
  dplyr::filter(gene %in% ENSG_code) %>%
  dplyr::select(-c(gene, cohort)) %>% 
  dplyr::select(c(genename, tissue, n.snps.in.data, 
                  n.snps.in.model, pred.perf.R2))

DT::datatable(A4_table, 
              rownames = FALSE,
              filter = 'top',
              extensions = 'Buttons',
              options = list(pageLength = 10,
                             dom = 'Bfrtip',
    buttons = c('copy', 'csv'))) %>% 
      DT::formatRound(columns = c("pred.perf.R2"), digits = 4)
```

###### 3.1.2.2 ACT


```
ACT_table <-
  readr::read_csv(
    paste0(path, 
    "input/PrediXcanTissueTables/ACT_predicted_expression_qc.csv")
  )

ACT_table <- ACT_table %>%
  dplyr::filter(gene %in% ENSG_code) %>%
  dplyr::select(-c(gene, cohort)) %>%
    dplyr::select(c(genename, tissue, n.snps.in.data, 
                  n.snps.in.model, pred.perf.R2))

DT::datatable(ACT_table, 
              rownames = FALSE,
              filter = 'top',
              extensions = 'Buttons',
              options = list(pageLength = 10,
                             dom = 'Bfrtip',
    buttons = c('copy', 'csv'))) %>% 
      DT::formatRound(columns = c("pred.perf.R2"), digits = 4)
```

###### 3.1.2.3 ADNI


```
ADNI_table <-
  readr::read_csv(
    paste0(path, "input/PrediXcanTissueTables/ADNI_predicted_expression_qc.csv")
  )

ADNI_table <- ADNI_table %>%
  dplyr::filter(gene %in% ENSG_code) %>%
  dplyr::select(-c(gene, cohort)) %>% 
    dplyr::select(c(genename, tissue, n.snps.in.data, 
                  n.snps.in.model, pred.perf.R2))

DT::datatable(ADNI_table,
              rownames = FALSE,
              filter = 'top',
              extensions = 'Buttons',
              options = list(pageLength = 10,
                             dom = 'Bfrtip',
    buttons = c('copy', 'csv'))) %>% 
      DT::formatRound(columns = c("pred.perf.R2"), digits = 4)
```

###### 3.1.2.4 ROS/MAP


```
ROSMAP_table <-
  readr::read_csv(
    paste0(path, 
    "input/PrediXcanTissueTables/ROSMAP_predicted_expression_qc.csv")
  )

ROSMAP_table <- ROSMAP_table %>%
  dplyr::filter(gene %in% ENSG_code) %>%
  dplyr::select(-c(gene, cohort)) %>% 
    dplyr::select(c(genename, tissue, n.snps.in.data, 
                  n.snps.in.model, pred.perf.R2))

DT::datatable(ROSMAP_table, 
              rownames = FALSE,
              filter = 'top',
              extensions = 'Buttons',
              options = list(pageLength = 10,
                             dom = 'Bfrtip',
    buttons = c('copy', 'csv'))) %>% 
    DT::formatRound(columns = c("pred.perf.R2"), digits = 4)
```


```
rm(A4_table)
rm(ACT_table)
rm(ROSMAP_table)
rm(ADNI_table)
```

##### 3.1.3 Meta Analysis


```
# combine table with globalres and globalres nc 
# read in meta analysis data
meta_analysis_results <-
  readr::read_csv(
    paste0(path, "input/prediXcanMetaAnalysis/combine/meta_analysis_results.csv")
  )

# filter for gene of interest
meta_analysis_results <- meta_analysis_results %>%
  dplyr::filter(ENSG %in% ENSG_code) 

# select specific columns
meta_analysis_results <- meta_analysis_results %>%
  dplyr::select(Gene, Tissue, resilience_type, beta, se, pvalue)

# table
DT::datatable(meta_analysis_results,
              rownames = FALSE,
              filter = 'top',
              extensions = 'Buttons',
              options = list(pageLength = 10,
                             dom = 'Bfrtip',
    buttons = c('copy', 'csv'))) %>%
  DT::formatRound(columns = c("beta", "se", "pvalue"), digits = 3) %>% 
  DT::formatStyle(c("pvalue"),
                  color = DT::styleInterval(0.05, c("red", "black")))
```

##### 3.1.4 GBJ on Meta Analysis


```
# read in meta analysis data
GBJ_meta_analysis <-
  readr::read_csv(
    paste0(path, "input/Predixcan_GBJ_meta_analysis.csv")
  )
GBJ_meta_analysis <- data.frame(GBJ_meta_analysis)
# filter for specific gene of interest
GBJ_meta <- GBJ_meta_analysis %>%
  dplyr::filter(ENSG %in% ENSG_code) %>% 
  dplyr::select(-c(ENSG))

GBJ_meta <-
  format(GBJ_meta, digits = 3) # change decimals

# table
DT::datatable(GBJ_meta,
              rownames = FALSE,
              extensions = 'Buttons',
              options = list(
                             dom = 'Bfrtip',
    buttons = c('copy', 'csv'))) %>%
  DT::formatRound(columns = c("GBJ", "GBJ_pvalue"), digits = 3) %>% 
  DT::formatStyle(c("GBJ_pvalue"),
                  color = DT::styleInterval(0.05, c("red", "black")))
```

#### 3.2 Predicted Expression - AD

##### 3.2.1 method

The following analysis result was pulled from research published in
this article: https://www.nature.com/articles/s41398-021-01677-0


```
igap_ad <- read_xls(paste0(path, "input/Predixcan_IGAP_2021.xls"))
# N = 65,534 obs of 17 variables
igap_ad <- data.frame(igap_ad)
```

##### 3.2.2 Tissue Table


```
ad_table <- igap_ad %>%
  dplyr::filter(symbol %in% gene_name) %>%
  dplyr::select(c(symbol, tissue, 
                  n_snps_used, n_snps_in_model, 
                  pred_perf_r2))
# N = 13

DT::datatable(ad_table, 
              rownames = FALSE,
              filter = 'top',
              extensions = 'Buttons',
              options = list(pageLength = 10,
                             dom = 'Bfrtip',
    buttons = c('copy', 'csv'))) %>% 
      DT::formatRound(columns = c("pred_perf_r2"), digits = 4)
```

##### 3.2.3 Meta Analysis


```
# do FDR correction
# FDR calculation
igap_ad <- igap_ad[order(igap_ad$pvalue),]
igap_ad$p.fdr <- p.adjust(igap_ad$pvalue, method = "fdr")

# filter for specific gene
meta_analysis_ad <- igap_ad %>%
  dplyr::filter(symbol %in% gene_name) %>%
  dplyr::select(c(symbol, tissue, 
                  zscore, pvalue, p.fdr))
# N = 13 
# attempts to try to get a better format for the decimals in significant figures
#meta_analysis_ad$pvalue <- signif(meta_analysis_ad$pvalue, digits = 3)
#meta_analysis_ad$p.fdr <- signif(meta_analysis_ad$p.fdr, digits = 3)
#meta_analysis_ad <- formatC(meta_analysis_ad$zscore, format = "e")  
#meta_analysis_ad <- formatC(meta_analysis_ad$pvalue, format = "e") 
#meta_analysis_ad <- formatC(meta_analysis_ad$p.fdr, format = "e") 

# table
DT::datatable(meta_analysis_ad,
              rownames = FALSE,
              filter = 'top',
              extensions = 'Buttons',
              options = list(pageLength = 10,
                             dom = 'Bfrtip',
    buttons = c('copy', 'csv'))) %>%
  DT::formatRound(columns = c("zscore", "pvalue", "p.fdr"), digits = 3) %>% 
  DT::formatSignif(columns = c("pvalue", "p.fdr"), digits = 3) %>% 
  DT::formatStyle(c("p.fdr"),
                  color = DT::styleInterval(0.05, c("red", "black")))
```
